# A novel, dynein-independent mechanism focuses the endoplasmic reticulum around spindle poles in dividing *Drosophila* spermatocytes

**DOI:** 10.1101/574855

**Authors:** Darya Karabasheva, Jeremy T. Smyth

**Affiliations:** Department of Anatomy, Physiology, and Genetics, Uniformed Services University of the Health Sciences, F. Edward Hébert School of Medicine, Bethesda, MD 20814

## Abstract

In dividing animal cells the endoplasmic reticulum (ER) concentrates around the poles of the spindle apparatus by associating with astral microtubules (MTs), and this association is essential for proper ER partitioning to progeny cells. The mechanisms that associate the ER with astral MTs are unknown. Because astral MT minus-ends are anchored by centrosomes at spindle poles, we tested the hypothesis that the MT minus-end motor dynein mediates ER concentration around spindle poles. Live *in vivo* imaging of *Drosophila* spermatocytes undergoing the first meiotic division revealed that dynein is required for ER concentration around centrosomes during interphase. In marked contrast, however, dynein suppression had no effect on ER association with astral MTs and concentration around spindle poles in early M-phase. Importantly though, there was a sudden onset of ER-astral MT association in *Dhc64C* RNAi cells, revealing activation of an M-phase specific mechanism. ER redistribution to spindle poles also did not require non-claret disjunctional (ncd), the other known *Drosophila* MT minus-end motor, nor Klp61F, a MT plus-end motor that generates spindle poleward forces. Collectively, our results suggest that a novel, M-phase specific mechanism of ER-MT association that is independent of MT minus-end motors is required for proper ER partitioning in dividing cells.

## Introduction

The endoplasmic reticulum (ER) cannot be formed by cells *de novo* and must be inherited during the process of cell division (Du et al., 2004). While the essential roles of the ER in the biogenesis of proteins, lipids and steroid hormones, as well as calcium signaling, are well recognized, little is known about the molecular mechanisms that ensure proper partitioning of the ER to progeny cells. This knowledge is fundamental to understanding the role of the ER in cell division and stem cell biology, with important implications for proper development, tissue repair, and cancer. Specifically, recent findings indicate that asymmetric partitioning of the ER, and misfolded proteins that accumulate there, has a critical role in maintaining pluripotency in stem cells as they undergo rapid cycles of cell division (Eritano et al., 2017; Moore et al., 2015; Smyth et al., 2015). Related evidence that ER functions contribute to the proliferative capacity and drug resistance of cancer cells is under active investigation, with an objective of revealing novel therapeutic strategies for treating malignancies (Avril et al., 2017; Urra et al., 2016). The overall goal of the present study was to establish new understanding of the cellular mechanisms that guide the localization of the ER during cell division, bringing us closer to full comprehension of the essential roles of the ER in normal physiology and disease.

Recent data suggest that proper partitioning of the ER during cell division, or M-phase, depends on specific association of the organelle with astral microtubules (MTs) of the mitotic spindle in both symmetrically and asymmetrically dividing cells (Smyth 2015). However, the specific factors that link the ER to astral MTs remain unknown. Identification of these factors is therefore an important next step in understanding mitotic ER partitioning. Most of our knowledge of ER-MT associations comes from non-dividing interphase cells, in which the ER is distributed throughout the cytoplasm with a distinct clustering or focus around centrosomes, the major MT organizing centers of cells. This distribution depends on MT motor-dependent movements of the ER toward both MT plus-ends and minus-ends, as well as stable attachments of the ER along MT filaments mediated by ER membrane proteins including REEPs, spastin, and CLIMP-63 (Gurel et al., 2014). Transport of the ER toward MT minus-ends is mediated by dynein motors, which are also responsible for focusing the ER around centrosomes where MT minus-ends are anchored (Wang et al., 2013; Wozniak et al., 2009). Conversely, MT plus-end transport is likely mediated by kinesins (Wozniak et al., 2009), and also depends on association with growing MT tips mediated, at least in part, by ER membrane embedded STIM1 and STIM2 proteins (Grigoriev et al., 2008). Collectively these associations point to a carefully orchestrated interplay between MT plus-end and minus-end directed transport mechanisms that determine the cellular distribution of the ER.

During the course of M-phase, there is a dramatic reorganization of the MT cytoskeleton, whereby the spindle apparatus forms with MT minus-ends anchored at the spindle poles by centrosomes. As this occurs, the majority of the ER becomes focused around the two centrosomes at the spindle poles and along astral MTs, and virtually none is found in the kinetochore region of the spindle where MT plus-ends reside (Schlaitz, 2014). Thus, there appears to be a shift from the balanced MT plus- and minus-end directed ER distribution during interphase to predominantly minus-end directed localization around spindle poles during M-phase. Consistent with this conclusion, tracking of the ER with growing MT plus-ends is inhibited during cell division due to mitosis-specific phosphorylation of STIM1 (Smyth et al., 2012; Smyth et al., 2009). This suppression of MT plus-end directed ER transport, and the totality of ER distribution toward MT minus-ends around spindle poles, suggests a predominant role for the MT-minus end motor dynein in M-phase specific ER distribution. Notably, dynein is highly localized to astral MTs and spindle poles in dividing cells (Busson et al., 1998; Quintyne and Schroer, 2002) and is required for the spindle pole localization of endosomes (Hehnly and Doxsey, 2014). However, despite this compelling case, a definitive role for dynein in M-phase ER distribution has never been directly established. Determining dynein’s role in the dramatic redistribution of the ER to spindle poles in M-phase is therefore important to understanding the mechanisms that ensure proper ER partitioning to progeny cells (Smyth et al., 2015)

The primary goal of this study was to test the hypothesis that dynein is required for astral MT association and spindle pole focusing of the ER in dividing cells. We accomplished this by live timelapse imaging of *Drosophila* spermatocytes undergoing the first meiotic division of spermatogenesis *in vivo*. This experimental system is particularly well suited for this investigation, as *Drosophila* spermatocytes allow for the analysis of dividing cells in a physiologic environment and have contributed greatly to our understanding of fundamental cell division mechanisms including spindle formation and cytokinesis (Giansanti and Fuller, 2012). Moreover, *Drosophila* spermatocytes are large cells that exhibit clearly defined redistribution of the ER onto astral MTs early in meiosis (Smyth et al., 2015; Tates, 1971). Here we present the surprising finding that although dynein is required for peri-centrosomal focusing of the ER during interphase, it does not mediate the astral MT-dependent spindle pole focusing of the ER around centrosomes during M-phase. Surprisingly, our results further reveal that redistribution of the ER toward MT minus-ends in dividing cells is mediated by a mechanism of ER-MT association that is entirely specific to M-phase and does not operate during interphase. This report lays the groundwork for identification of this novel mechanism of ER-MT association that is essential for the process of ER inheritance.

## Results

### Dynein mediates ER association with centrosomal MTs during interphase

We began our investigation by analyzing the role of dynein in ER distribution throughout the cell cycle in *Drosophila* spermatocytes. *Drosophila* spermatocytes are organized in cysts of 16 cells, and these cells synchronously undergo two meiotic divisions to form 64 haploid spermatids (Fuller, 1993). During late interphase prior to the first meiotic division, spermatocytes have two cortically located centrosomes with elaborate MT asters. Double labeling of ER and MTs by co-expression of ER-targeted RFP (RFP-KDEL) and GFP-tubulin, respectively, revealed that the ER was widely distributed throughout the cytoplasm during interphase, but also exhibited a distinct concentration around each cortical centrosome, with segments of the ER closely aligned along the centrosomal MTs (Figure 1a). The ratio of RFP-KDEL fluorescence intensity immediately adjacent to each centrosome versus that elsewhere in the cytoplasm (ER_centrosome_ / ER_cytoplasm_) was 1.60 ± 0.06 (mean ± SEM; n = 5 cells) in interphase cells, reflecting 60% enrichment of the ER around centrosomes as compared to the rest of the cytoplasm at this cell cycle stage. We also observed clear concentration of the ER around centrosomes using GFP labeled protein disulfide isomerase (GFP-PDI) to label the organelle and ana1-tdTomato to identify centrosomes, demonstrating that the observed distribution of ER is not ER label specific (Figure 1b). To test whether this peri-centrosomal ER concentration in interphase is mediated by dynein, we expressed inducible RNAi targeting *Drosophila* dynein heavy chain (D*hc64C*) specifically in spermatocytes using the germ cell-specific *bam-GAL4:VP16* driver (Chen and McKearin, 2003). This approach avoided the lethality observed in loss-of-function dynein mutant flies (Gepner et al., 1996), while allowing for the use of a highly effective interfering RNA strategy that reduced *Dhc64C* mRNA levels by 77% in whole larvae when driven with the ubiquitous *da-GAL4* driver (Supplementary Fig. S1). Strikingly, in *bam-GAL4:VP16*-driven *Dhc64C* RNAi spermatocytes the ER was entirely excluded from centrosomal MTs, resulting in complete absence of the peri-centrosomal concentration of ER seen in controls (Figure 1a). This resulted in an ER_centrosome_ / ER_cytoplasm_ ratio of 0.27 ± 0.02 (mean ± SEM; n = 5 cells) in *Dhc64C* RNAi spermatocytes, a statistically significant difference from controls (see Figure 2d) and a clear representation of the observed absence of ER around the centrosomes as compared to the rest of the cytoplasm. These effects of dynein suppression on interphase ER distribution were again recapitulated with the GFP-PDI probe (Figure 1b). Together, these results indicated that dynein mediates the association of the ER with MTs, and functions to focus the ER around MT minus-ends at centrosomes in interphase *Drosophila* spermatocytes.

**Figure 1.**
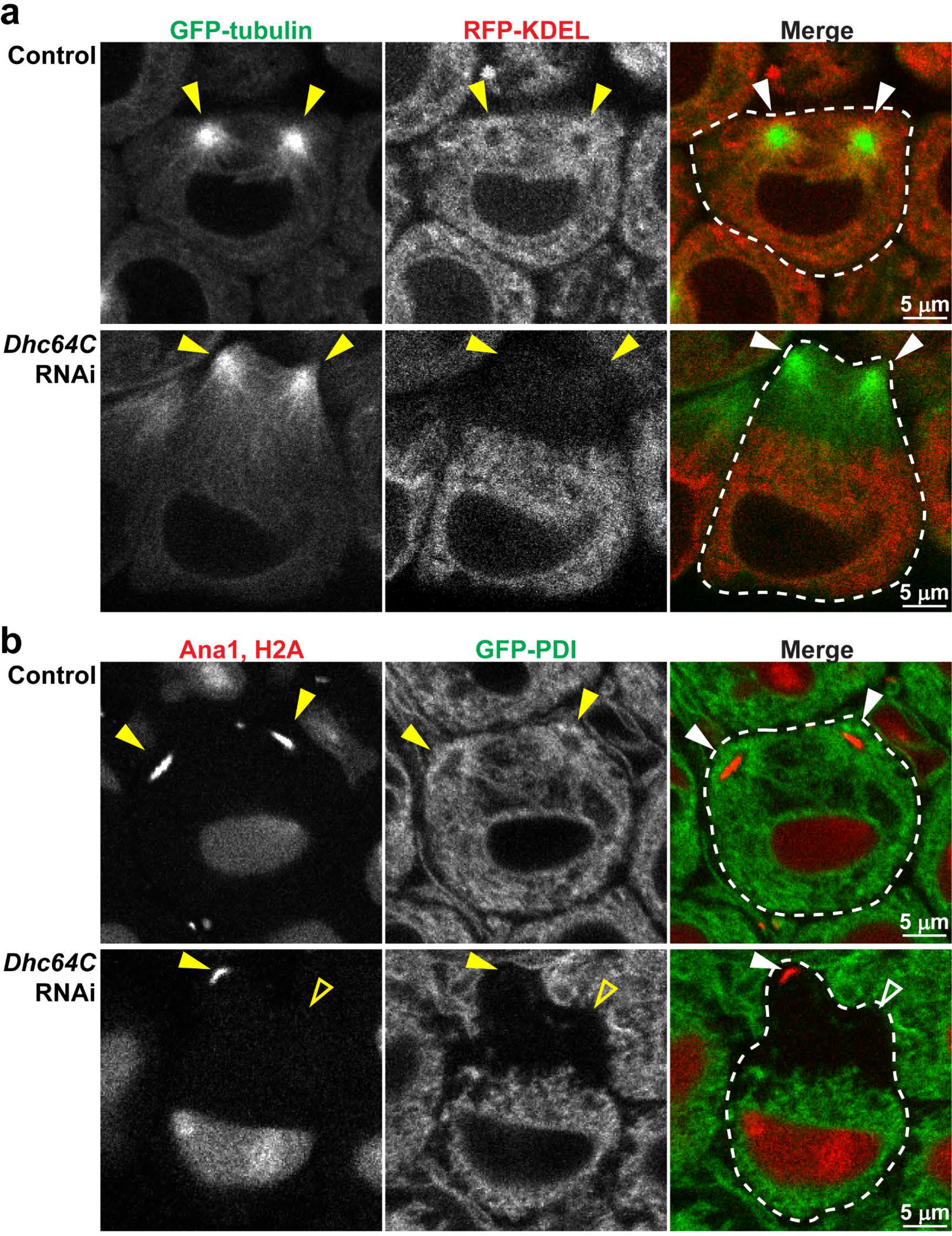
Dynein is required for ER concentration around centrosomes in interphase. a) Representative images of control (upper panel) and *Dhc64C* RNAi (lower panel) spermatocytes co-expressing GFP-tubulin (green) and RFP-KDEL (red) during late interphase. b) Representative images of control (upper panel) and *Dhc64C* RNAi (lower panel) spermatocytes co-expressing Ana1-tdTomato, H2A-RFP (red) and GFP-PDI (green) during late interphase. Dotted lines outline individual cells, and filled arrowheads point to visible centrosomes. Open arrowheads indicate the approximate location of centrosomes that are out of the focal plane.

**Figure 2.**
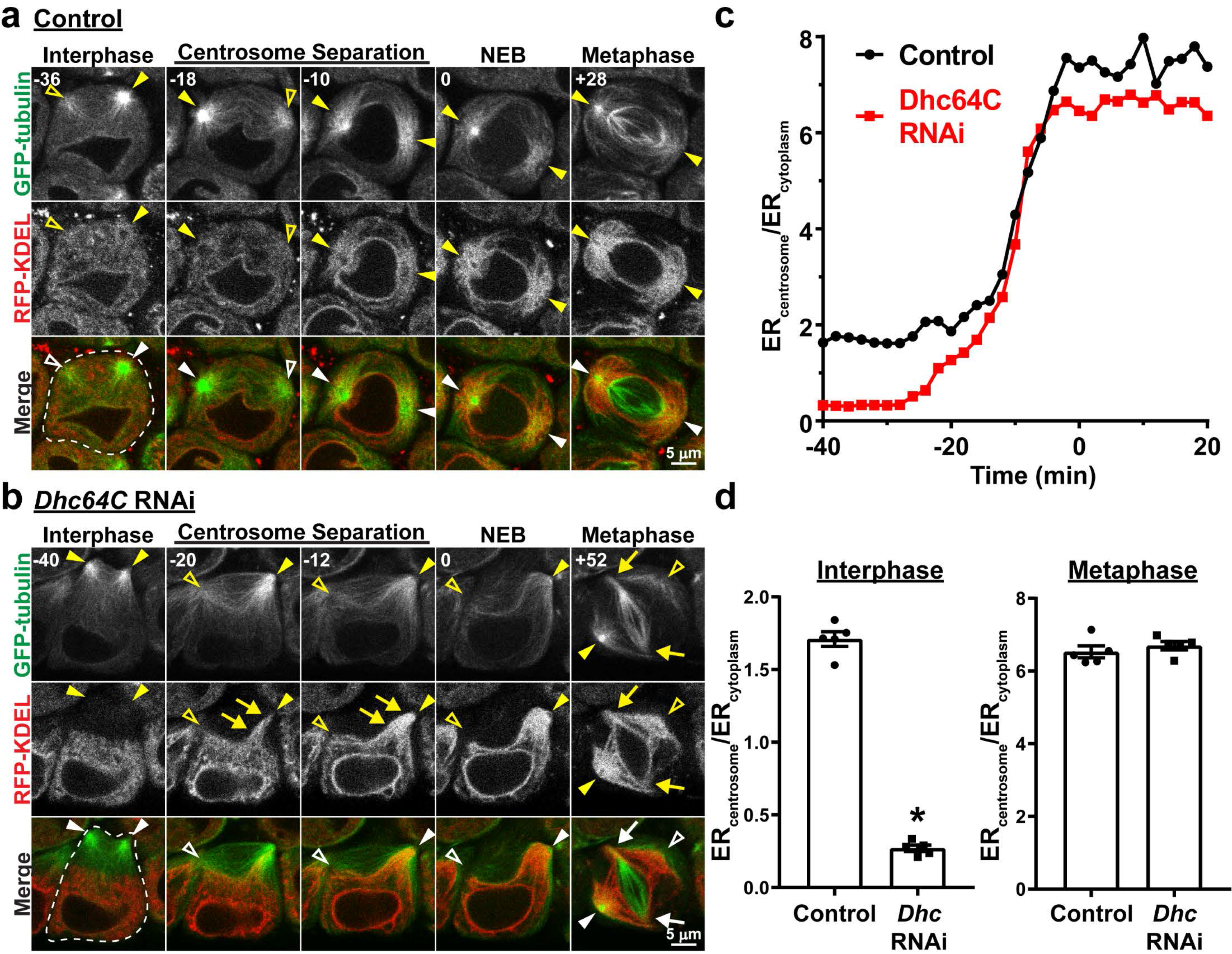
ER association with astral MTs during M-phase is independent of dynein activity. a) Representative images of a control spermatocyte co-expressing GFP-tubulin (green) and RFP-KDEL (red) from late interphase through metaphase of the first meiotic division. Times in minutes in the upper left of each image are reported relative to NEB, which was assigned as the first timepoint at which there was a notable increase in GFP-tubulin fluorescence in the nucleus. Filled arrowheads indicate visible centrosomes, and open arrowheads indicate the approximate location of centrosomes that are out of the focal plane. b) Representative images of a *Dhc64C* RNAi spermatocyte co-expressing GFP-tubulin (green) and RFP-KDEL (red) from late interphase through metaphase of the first meiotic division. Times in minutes in the upper left of each image are reported relative to NEB. Filled arrowheads indicate visible centrosomes, and open arrowheads indicate the approximate location of centrosomes that are out of the focal plane. Arrows in the “Centrosome Separation” images (−20 and −12 min) point to RFP-KDEL – labeled ER that is moving toward the centrosome along astral MTs. Arrows in the “Metaphase” image (+52 min) point to the spindle poles where the kinetochore MTs are focused; note that the centrosomes with astral MTs and associated ER are completely dissociated from the spindle poles. c) The ratio of RFP-KDEL fluorescence intensity immediately adjacent to a single centrosome versus that in the cytoplasm near the nucleus (ER_centrosome_ / ER_cytoplasm_) was calculated every 2 minutes from late interphase (−40 min) through metaphase (20 min) for the control and *Dhc64C* RNAi cells shown in a) and b), respectively. Time is relative to NEB. d) Mean ER_centrosome_ / ER_cytoplasm_ ± SEM for five control and *Dhc64C* RNAi cells during interphase (left panel; measured at 40 min prior to NEB) and metaphase (right panel; measured at 20 min post NEB). ‘*’, p < 0.0001, t-test.

### In contrast to interphase, dynein is not required for ER concentration around centrosomes at spindle poles during M-phase

During M-phase in *Drosophila* spermatocytes, the majority of the ER is associated with astral MTs around the two spindle pole centrosomes, and a fraction of ER membranes also envelopes the spindle apparatus and is referred to as the parafusorial membranes (Fuller, 1993; Tates, 1971) (see Figure 2a, “Metaphase”). We previously demonstrated that the specific association of ER with astral MTs is required for proper ER inheritance in dividing cells (Smyth 2015).

Accordingly, we next sought to determine whether this astral MT association of ER and concentration around spindle pole centrosomes in M-phase depends on dynein activity, similar to what we observed in interphase. As shown in Figures 2a, 3a, Supplementary Videos 1 and 2, the first visible event of M-phase in spermatocytes was separation and movement of the two centrosomes from the cell cortex to opposite sides of the nucleus, where they established the two poles of the meiotic spindle. As the centrosomes migrated to the nuclear envelope in control cells, there was a marked and steady increase in the amount of ER associated with the astral MTs around each centrosome, and a concomitant decrease in ER in the rest of the cytoplasm. At the stage of nuclear envelope breakdown (NEB) and continuing through metaphase, nearly the entire cellular content of ER was associated with the astral MTs at the two spindle poles. This dramatic redistribution of the ER from the cytoplasm to astral MTs around the centrosomes was clearly reflected by a four-fold increase in ER_centrosome_ / ER_cytoplasm_ at metaphase as compared to interphase (6.45 ± 0.26 at metaphase vs. 1.60 ± 0.06 at interphase; Figure 1c,d). In *Dhc64C* RNAi spermatocytes, the centrosomes separated but they failed to move from the cell cortex to the nuclear envelope, consistent with dynein’s role in centrosome-nuclear envelope engagement (Robinson et al., 1999; Splinter et al., 2010; Wainman et al., 2009). Despite this clear and specific effect of dynein suppression on the movement of centrosomes, in these same cells the ER exhibited a sudden and rapid movement onto the astral MTs and toward the cortical centrosomes similar to that observed in control spermatocytes (Figure 2b, 3b, Supplementary Videos 3 and 4). Close temporal examination revealed that the ER moved processively toward the cortical centrosomes in *Dhc64C* RNAi spermatocytes, resulting in concentration of the ER around the centrosomes by the time of NEB (Figure 3b’). This movement of the ER toward the centrosomes in dynein suppressed cells was nevertheless MT-dependent, because it was inhibited by the MT depolymerizing drug colchicine (Figure 4a,b). Importantly, while the redistribution of ER to centrosomes during M-phase was not affected by *Dhc64C* RNAi, spindle architecture was greatly altered as expected with dynein suppression. Specifically, as *Dhc64C* RNAi cells progressed into a metaphase-like stage with congressed chromosomes and organized kinetochore spindle fibers, the centrosomes with their astral MTs were dissociated from the spindle poles, consistent with dynein suppression (Morales-Mulia and Scholey, 2005; Wainman et al., 2009); in some cases the axis of the two centrosomes was completely perpendicular to the orientation of the spindle itself (Figure 2b). Even with these dramatic defects in spindle architecture in *Dhc64C* RNAi spermatocytes, the ER remained specifically associated with the astral MTs around the centrosomes, as in control cells, suggesting that dynein is not required for M-phase redistribution of the ER to astral MTs around centrosomes. Further supporting this conclusion, the maximal ER_centrosome_ / ER_cytoplasm_ value attained in *Dhc64C* RNAi cells at metaphase was not significantly different from controls (Figure 2c,d). RNAi-mediated suppression of *NudE*, a dynein activator with multiple mitotic functions (Wainman et al., 2009), was used to further confirm the dynein-independence of ER redistribution to astral MTs in M-phase. To this end, the effects of *NudE* suppression were nearly identical to those of *Dhc64C*, on both ER redistribution and spindle architecture (Supplementary Fig. S2). Collectively, these results indicate that even though ER concentration around centrosomes is dynein dependent during interphase, redistribution of the ER onto astral MTs around spindle pole centrosomes during M-phase is independent of dynein activity. Further, the sudden onset of ER association with astral MTs in dynein-suppressed cells suggests that an M-phase specific mechanism of ER-MT association is engaged early in cell division to concentrate the ER around centrosomes.

**Figure 3.**
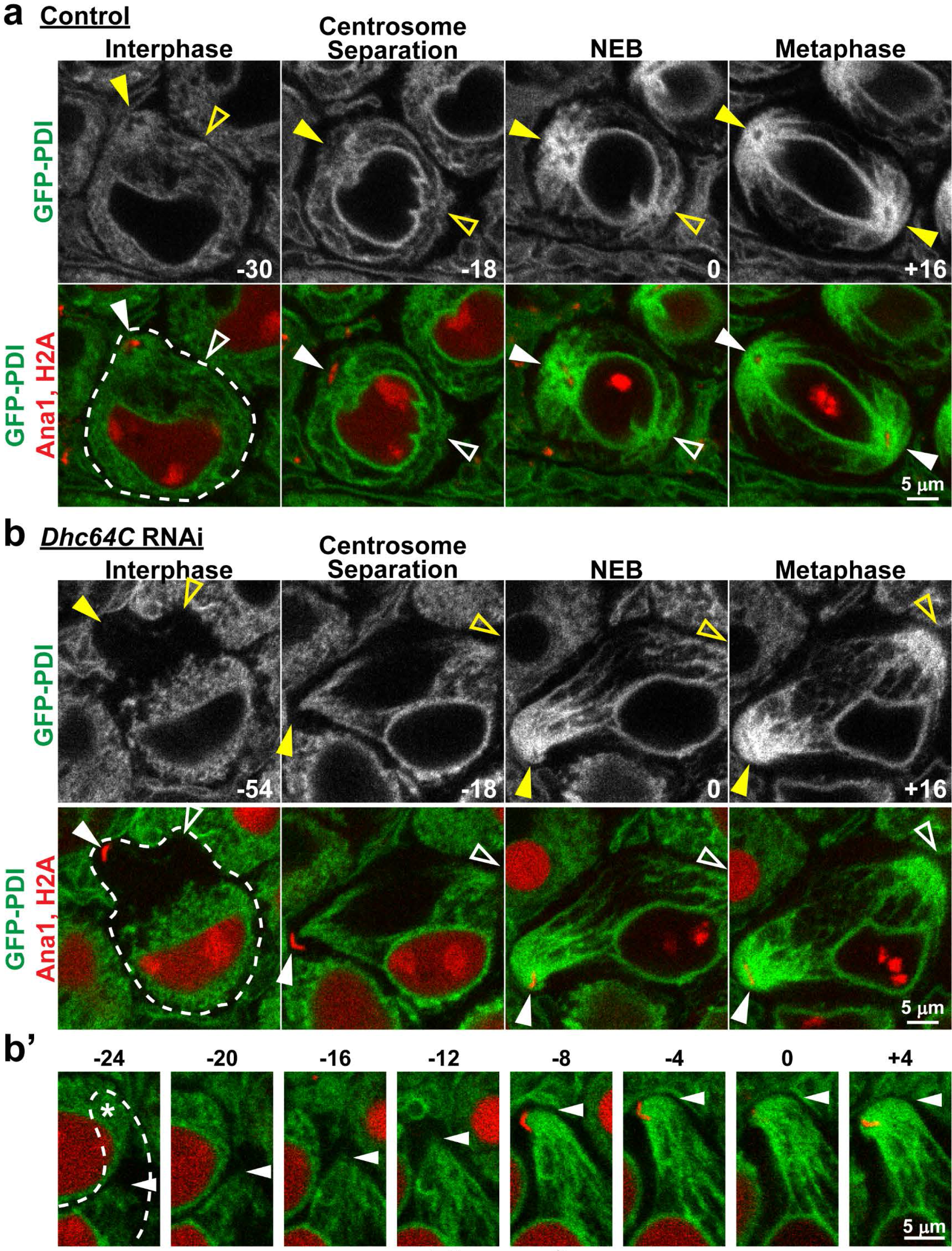
Spindle pole focusing of the ER during M-phase is independent of dynein activity. a) Representative images of a control spermatocyte co-expressing Ana1-tdTomato, H2A-RFP (red) and GFP-PDI (green) from late interphase through metaphase of the first meiotic division. Times in minutes are reported relative to NEB, which was assigned as the first timepoint at which the diffuse H2A-RFP fluorescence in the nucleus dissipated. Filled arrowheads indicate visible centrosomes marked by Ana1-tdTomato, and open arrowheads indicate the approximate location of centrosomes that are out of the focal plane. b) Representative images of a *Dhc64C* RNAi spermatocyte co-expressing Ana1-tdTomato, H2A-RFP (red) and GFP-PDI (green) from late interphase through metaphase of the first meiotic division. Times in minutes are reported relative to NEB. Filled arrowheads indicate visible centrosomes, and open arrowheads indicate the approximate location of centrosomes that are out of the focal plane. b’) Expanded timecourse of GFP-PDI being drawn toward one of the centrosomes in the *Dhc64C* RNAi spermatocyte. The arrowheads indicate the leading front of GFP-PDI. In the first four frames the centrosome to which GFP-PDI is being drawn is out of the plane of focus, but its approximate location is indicated by ‘*’ in the first frame.

**Figure 4.**
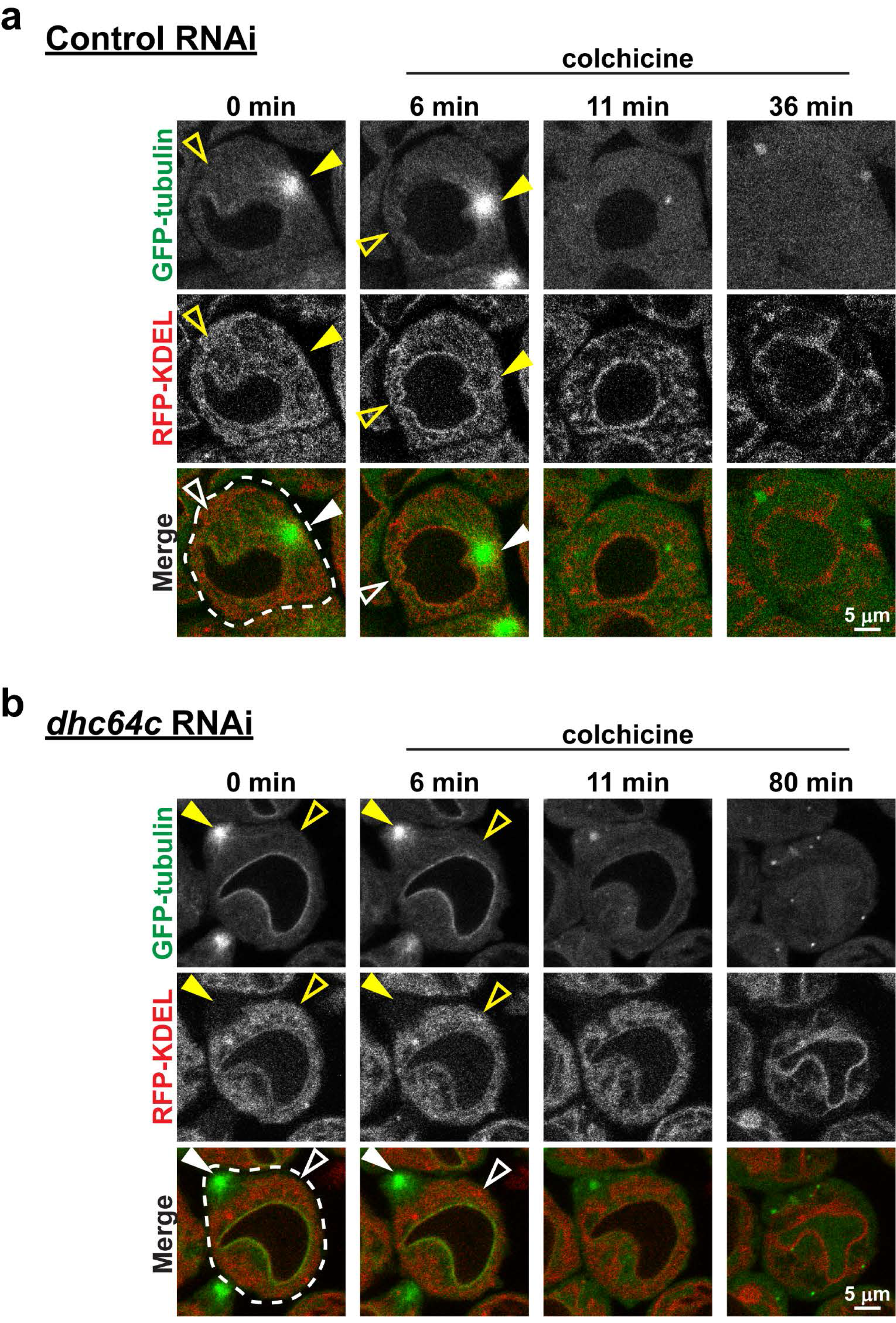
Dynein independent ER redistribution to centrosomes requires MTs. Control (a) and *Dhc64C* (b) RNAi spermatocytes expressing GFP-tubulin (green) and RFP-KDEL (red) were imaged beginning in late interphase before the commencement of ER redistribution to the centrosomes. Colchicine (50 μM) was then applied 6 min later and the cells were imaged through NEB to encompass the time during which ER redistribution would normally occur. Note the lack of any distinct focus of the ER by the time of NEB in the control cell, and that the ER fails to be drawn to the cell cortex, where the centrosome was located prior to colchicine application, in the *Dhc64C* RNAi cell.

### Neither ncd nor Klp61F mediate ER association with astral MTs during M-phase

The processive, MT-dependent movement of the ER toward centrosomes in dynein suppressed cells suggests that a different MT minus-end motor may be responsible for spindle pole concentration of the organelle. We therefore tested non-claret disjunctional (ncd; human kinesin-14 orthologue), the only other known MT minus-end motor in *Drosophila*, for its possible role in M-phase specific ER redistribution. Spermatocyte specific expression of an *ncd*-targeted RNAi construct that, when expressed globally resulted in approximately 55% knockdown of *ncd* transcript in whole larvae (Supplementary Fig. S1), had no observable effect on the concentration of ER around centrosomes during interphase as compared to controls (Figure 5a). Further, early spindle formation proceeded normally in *ncd* RNAi cells, and the most notable spindle phenotype observed was a deflection of interpolar MTs to one side in late metaphase, just prior to anaphase onset (Supplementary Fig. S3). Importantly, as seen in control and dynein suppressed cells, the ER redistributed to astral MTs beginning with centrosome migration in *ncd* RNAi cells (Figure 5a). Fluorescence intensity analysis confirmed these observations, with ER_centrosome_ / ER_cytoplasm_ values of 1.65 ± 0.03 and 6.55 ± 0.22 (mean ± SEM; n = 5 cells) for interphase and M-phase respectively, which were not significantly different from controls (Supplementary Fig. S4). To confirm that the lack of effects on ER redistribution as well as spindle architecture were not due to insufficient RNAi mediated knockdown of *ncd* expression, we also tested a hypomorphic *ncd* allele (*ncd*^*D*^) that produces ncd protein that lacks motor function (Endow et al., 1994; Komma et al., 1991). Similar to RNAi cells, spermatocytes from *ncd*^*D*^ homozygous larvae also formed near-normal spindles, and while these cells did exhibit a slight deflection of interpolar MTs just prior to anaphase onset, ER redistribution to astral MTs was not perturbed (Supplementary Fig. S3). We further considered the possibility that ncd and dynein may have complimentary functions in ER-astral MT association. To test this, we suppressed the expression of both D*hc64C* and *ncd* in spermatocytes by RNAi. The phenotype associated with combined dynein and ncd suppression was nearly identical to that observed with D*hc64C* RNAi alone: these cells had cortical centrosomes that failed to migrate to the NE, and the ER was excluded from the centrosomal MTs during interphase but was abruptly pulled toward the cortical centrosomes and onto astral MTs as early meiosis progressed (Figure 5b). This ER redistribution was also quantitatively similar to that in D*hc64C* alone suppressed cells as determined by ER_centrosome_ / ER_cytoplasm_ measurement (Supplementary Fig. S4). Taken together, these results suggest that neither ncd nor dynein is required for M-phase specific ER redistribution to astral MTs.

**Figure 5.**
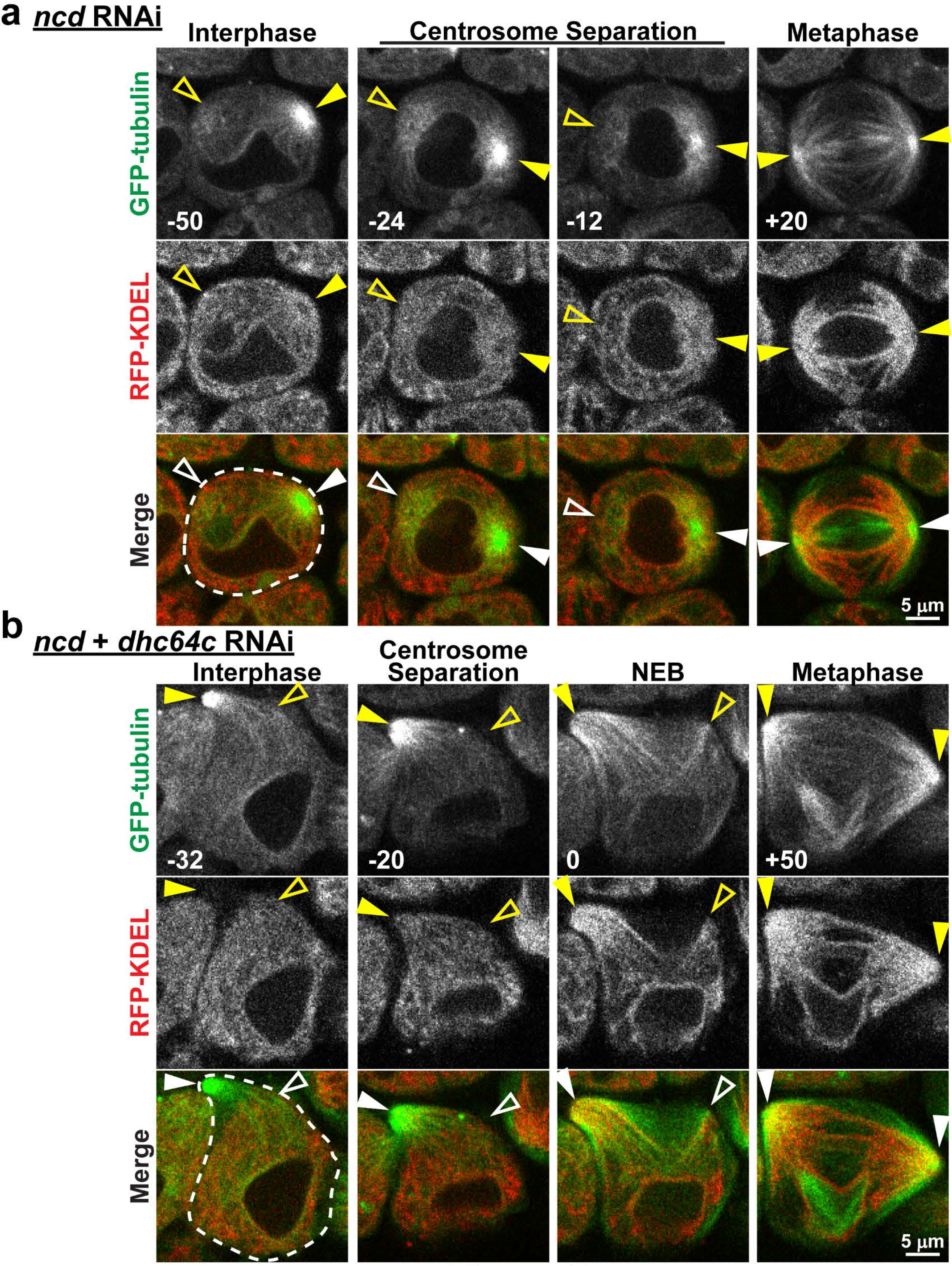
Ncd is not required for astral MT association of the ER in M-phase. a) Representative images of a *ncd* RNAi expressing spermatocyte with GFP-tubulin (green) and RFP-KDEL (red) from late interphase (−50 min) through metaphase (+20 min) of meiosis (times relative to NEB, lower left of images). Note the clear association of ER with astral MTs at metaphase (compare to Figure 1A). b) Representative images of a spermatocyte co-expressing *ncd* and *Dhc64C* RNAi, with GFP-tubulin (green) and RFP-KDEL (red), from late interphase (− 32 min) through metaphase (+50 min) of meiosis. Note the similarities to cells with *Dhc64C* RNAi alone (compare to Figure 2B). Filled arrowheads indicate visible centrosomes, and open arrowheads indicate the approximate location of centrosomes that are out of the focal plane.

Lastly, we considered the possibility that Klp61F (human kinesin-5 orthologue) may be involved in MT association and spindle pole focusing of the ER during M-phase. Klp61F is a MT plus-end kinesin motor that slides antiparallel, interpolar MTs, generating outward forces within the spindle that maintain spindle pole separation (Mann and Wadsworth, 2019). Thus, while Klp61F is not a *bona fide* MT minus-end motor, the spindle pole-directed forces that it generates could play a role in driving the ER toward spindle poles. *Klp61F* RNAi that resulted in approximately 76% knockdown of *Klp61F* transcript when expressed ubiquitously (Supplementary Fig. S1) had no effect on the early events of spindle assembly in spermatocytes; however, shortly after NEB the spindles collapsed into monopoles (Figure 6a) confirming the suppression of Klp61F activity by RNAi. Despite this defect in spindle architecture seen in Klp61F suppressed spermatocytes, the ER still correctly redistributed to astral MTs in early meiosis (Figure 6a), and the overall extent of redistribution of the ER to spindle poles was similar to controls based on ER_centrosome_ / ER_cytoplasm_ measurements (Supplementary Fig. S4). Notably, the ER remained focused around the spindle poles even as the spindle collapsed and the poles moved toward one another. We also tested *Klp61F* in combination with *Dhc64C* suppression, but again the ER was drawn along astral MTs to the cortical centrosomes in early meiosis similar to cells with *Dhc64C* RNAi alone (Figure 6b, Supplementary Fig. S4). Collectively, these results reveal the existence of a novel, M-phase specific mechanism that mediates redistribution of ER to astral MTs in dividing cells.

**Figure 6.**
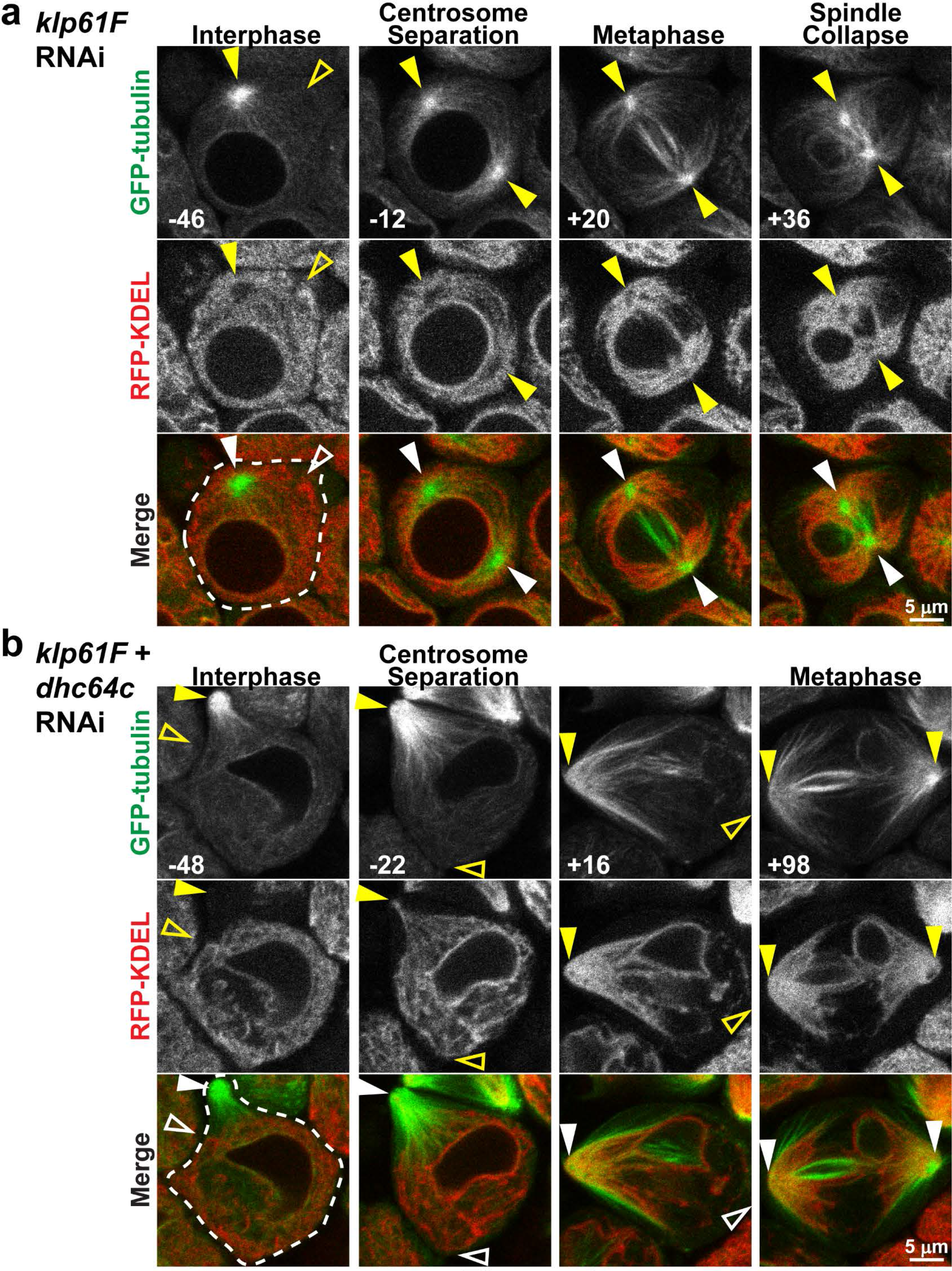
Klp61F is not required for astral MT association of the ER in M-phase. a) Representative images of a *Klp61F* RNAi expressing spermatocyte with GFP-tubulin (green) and RFP-KDEL (red) from late interphase (−46 min) through metaphase (+36 min) of meiosis (times relative to NEB, lower left of images). Note the collapse of the spindle with movement of the two spindle poles toward one another during metaphase (+36 min). b) Representative images of a spermatocyte co-expressing *Klp61F* and D*hc64c* RNAi with GFP-tubulin (green) and RFP-KDEL (red) from late interphase (−48 min) through metaphase (+98 min) of meiosis. Note the similarities to D*hc64C* RNAi alone (compare to Figure 2b). Filled arrowheads indicate visible centrosomes, and open arrowheads indicate the approximate location of centrosomes that are out of the focal plane.

## Discussion

Proper distribution of the ER in dividing cells is critical, because it ensures that progeny cells receive necessary proportions of this essential organelle. Disruptions in this process may result in cells that are prone to accumulation of misfolded proteins and ER stress, as well as dysregulation of lipid homeostasis and calcium signaling, all of which are involved in a spectrum of diseases including cancer, neurodegeneration, and diabetes. However, the importance of mitotic ER distribution may extend beyond organelle inheritance. For example, it was recently demonstrated that the ER plays an essential role in restricting the distribution of damaged proteins in asymmetrically dividing stem cells. This mechanism, which depends on proper distribution of the ER around the mitotic spindle, may allow vital stem cells to protect themselves by asymmetrically shuttling damaged proteins to non-essential progeny cells (Moore et al., 2015). In addition, it has been suggested that the ER delivers highly localized calcium signals that are essential for proper function of the spindle apparatus (Whitaker, 2006). Thus, disruption of ER distribution around the mitotic spindle may impair the process of cell division itself with potentially devastating effects on development and tissue homeostasis. Conversely, therapeutic interventions that specifically target ER functions in cancer cells may prove to be effective treatments in neoplastic disease (Banerjee and Zhang, 2018). Identification of the molecular mechanisms that regulate M-phase specific distribution of the ER is critical to understanding these multi-faceted roles for the organelle in dividing cells. The present investigation demonstrates the existence of a novel mechanism that mediates ER-MT association in a manner that is specific to M-phase and independent of dynein and other known MT motors.

The first studies to examine ER distribution in dividing cells reported the organelle’s distinct organization around the poles of the mitotic spindle, where MT minus-ends are clustered (Harris, 1975; Waterman-Storer et al., 1993). Subsequent studies have confirmed this observation across multiple species, suggesting the existence of a unifying mechanism of ER partitioning that involves MT-dependent spindle pole association (Bobinnec et al., 2003; Smyth et al., 2015; Terasaki, 2000). Accordingly, involvement of a MT minus-end motor to move the ER toward spindle poles has been suggested in multiple studies (Bobinnec et al., 2003; Schlaitz et al., 2013; Waterman-Storer et al., 1993), with dynein being the most likely candidate. Importantly, dynein associates with ER-rich microsomal fractions (Allan, 1995; Lane et al., 2001; Steffen et al., 1997) and is required for proper ER distribution in interphase cells and extracts (Wang et al., 2013; Wedlich-Soldner et al., 2002; Wozniak et al., 2009), suggesting that the motor can associate with and transport ER membranes. Surprisingly however, the putative role for dynein in regulating ER distribution during cell division has not been directly tested. We therefore addressed this question using meiotic *Drosophila* spermatocytes, which demonstrate a dramatic, near complete MT-dependent redistribution of the ER to spindle poles during cell division. Our results clearly demonstrate a role for dynein in ER transport and localization in these cells, because ER distribution toward MT minus-ends and around centrosomes during interphase was completely disrupted by dynein suppression. Surprisingly, however, dynein was dispensable for astral MT association and spindle pole focusing of the ER during meiosis. Importantly and consistent with previous reports (Robinson et al., 1999; Splinter et al., 2010; Wainman et al., 2009), dynein suppression had dramatic effects on spindle formation and architecture in *Drosophila* spermatocytes including failure of centrosome migration to the nuclear envelope and complete dissociation of spindle poles from the focused kinetochore MT fibers. Nevertheless, despite these highly abnormal spindles, we observed that early in meiosis the ER was still drawn onto the astral MTs and towards the centrosomes, despite the centrosomes’ mislocalization at the cell cortex. Moreover, this redistribution of the ER occurred with the same kinetics and to the same extent as in control cells, suggesting that the mechanism of ER-astral MT association is completely independent of dynein. In this regard, data indicating that dynein association with membranous organelles is inhibited during M-phase of cell division (Niclas et al., 1996) suggests the existence of a dynamic mechanism that reciprocally regulates dynein association of the ER with MTs according to the stage of the cell cycle.

An important outcome of dynein suppression in spermatocytes is that it allowed a clear separation of interphase versus M-phase mechanisms of ER association with centrosomal MTs, wherein the ER was completely excluded from centrosomal MTs during interphase but suddenly moved along MTs toward the centrosomes early in M-phase. This reveals that an M-phase specific mechanism of ER-MT association that concentrates the ER around centrosomes is triggered at the onset of cell division, possibly due to specific cyclin / cyclin-dependent kinase activity (Bergman et al., 2015). This conclusion is further supported by the observations that several of the mechanisms for ER-MT association that operate during interphase are in fact inhibited during M-phase, including those mediated by STIM1 (Smyth et al., 2012), CLIMP-63 (Vedrenne et al., 2005), and possibly dynein (Niclas et al., 1996). Collectively, these findings suggest an enticing mechanism whereby most or all interphase mechanisms of ER-MT association are inhibited during M-phase, allowing an M-phase specific mechanism to predominate and ensure ER association with astral MTs and proper partitioning to daughter cells.

The challenge moving forward is to identify the M-phase specific mechanism that associates the ER with astral MTs. Importantly, in dynein suppressed spermatocytes it was clear that the ER moved along astral MTs toward the minus-ends, suggesting that a MT minus-end motor distinct from dynein may be involved. However, the present findings indicate that neither ncd, the only other known *bona fide* MT minus-end motor in *Drosophila*, nor Klp61F, which generates pole-ward forces in the spindle, are involved in the unique M-phase distribution of ER with astral MTs. Thus, it is possible that an unidentified MT minus-end motor is required, or that a non-motor factor stably attaches ER membranes along MT fibers and tubulin flux then moves these membranes toward centrosomes. In this regard, it was recently demonstrated that human REEP3 and 4, members of the REEP1-4 family of ER membrane proteins, directly associate with MTs and play a role in spindle pole focusing of the ER (Schlaitz et al., 2013). However, whether REEP3/4 associate the ER specifically with astral MTs has yet to be determined. *Drosophila* have a single orthologue to human REEPs1-4, known as *REEPA* (Yalcin et al., 2017), and our preliminary observations indicate that REEPA also is not required for astral MT association of the ER in meiotic *Drosophila* spermatocytes (unpublished observations). It is also important to note that different mechanisms of ER-MT association may operate during cell division in different cell types. Thus, while our results cannot rule out a role for dynein or other MT minus-end motors in ER partitioning in cells other than *Drosophila* spermatocytes, our findings do indicate that these factors are not universally required.

In addition to identifying the molecular factors that link the ER to astral MTs during cell division, another important question is how does this association discriminate astral MTs from other MTs of the spindle apparatus? Certainly, suppression of interphase mechanisms plays a role, as expression of nonphosphorylatable STIM1, which remains associated with MTs during mitosis, results in mislocalization of the ER to kinetochore MTs (Smyth et al., 2012). It is also possible that differences in tubulin post-translational modifications facilitate discrimination between different MT populations within the spindle, as demonstrated for kinesin-7 motors that carry chromosomes toward the spindle equator along detyrosinated MTs of the inner spindle (Barisic et al., 2015).

In conclusion, we have demonstrated that an M-phase specific mechanism associates the ER with astral MTs and partitions the organelle to spindle poles in dividing cells. Surprisingly, and in contrast to many previous suggestions, this ER localization is not mediated by dynein or other known MT minus-end motors. Identification of the mechanisms involved is an important next step in better understanding the role of the ER in cell division, stem cell longevity and pluripotency, and tissue architecture. This knowledge will bring us closer to developing novel therapeutics for pathological processes underlying cancer and age-related tissue degeneration.

## Materials and Methods

### Fly Stocks and Husbandry

The following fly stocks were obtained from the Bloomington *Drosophila* stock center: RFP-KDEL (Stock# 30910; expresses RFP targeted to the ER lumen under control of the *UASp* promoter); His2A-RFP (Stock# 23651; expresses Histone 2Av tagged with mRFP in all cells); GFP-PDI (Stock# 6839; expresses PDI tagged with GFP at the endogenous locus (Kelso et al., 2004)) *ncd*^*D*^(Stock# 2243); *Dhc64C* RNAi (Stock# 36698); *NudE* RNAi (Stock# 38954); and da-GAL4 (Stock# 55850; expresses GAL4 ubiquitously under control of the *daughterless* promoter). Lines carrying RNAi targeting *Klp61F* and *ncd* were obtained from the Vienna *Drosophila* Resource Center (Stock# 109280 and 110255, respectively). Stocks expressing GFP-tagged α-tubulin84B under control of the ubiquitin promoter (GFP-tubulin) (Rebollo et al., 2004), tdTomato-tagged Ana1 under control of the *UASp* promoter (Blachon et al., 2008), and Sec61α tagged with GFP at the endogenous promoter (Morin et al., 2001) were also used. The *bam-GAL4:VP16* driver line was a generous gift from Michael Busczcak. RFP-KDEL was recombined with GFP-tubulin using standard techniques for experiments in which both transgenes were expressed, as were His2A-RFP and ana1-tdTomato. Flies were maintained on standard cornmeal agar food, and all crosses were run at 25º C. Spermatocyte-specific RNAi experiments were carried out by crossing animals carrying *bam-GAL4:VP16* to animals carrying the respective *UASp*-regulated RNAi transgene. We attempted to use two control RNAi constructs available from the Bloomington Stock Center, one that targets *mCherry* and one that targets *GAL4*; however, both of these resulted in suppression of RFP-KDEL expression. Controls for RNAi experiments were therefore carried out by crossing RNAi animals to *w*^*1118.*^

### Live spermatocyte imaging

Whole testes were dissected from 3^rd^ instar larvae in *Drosophila* Schneider’s medium (Life Technologies) containing Antibiotic-Antomycotic (Life Technologies), and were mounted in the same medium for imaging on a 50 mm gas-permeable lumox dish (Sarstedt). The medium was surrounded by Halocarbon 700 oil (Sigma) to support a glass coverslip (22x22 mm, #1.5, Fisher) that was placed on top of the medium (Lerit et al., 2014). For colchicine treatments, whole testes were placed in a glass-bottom dish (MatTek) filled with Schneider’s medium. Colchicine was then added to a final concentration of 50 μM at the indicated times. Individual cysts of spermatocytes in whole testes were imaged with an inverted Nikon A1R scanning confocal microscope system using 488 nm laser excitation for GFP and 568 nm laser excitation for RFP and tdTomato through a 40x, 1.3 NA objective lens. Z-stacks compromising 15-20 images at 1 μm intervals were obtained every 1 to 2 minutes. All imaging was carried out at room temperature. Images were processed and videos were compiled using ImageJ software (NIH). All image times are reported relative to NEB in order to standardize each timecourse to the same event for each cell. NEB was determined by the first image in the timecourse in which GFP-tubulin fluorescence appeared in the nucleus or the diffuse RFP-H2A fluorescence in the nucleus diminished.

### ER_centrosome_ / ER_cytoplasm_ quantification

We defined ER_centrosome_ / ER_cytoplasm_ as the ratio of RFP-KDEL fluorescence intensity immediately adjacent to a single centrosome over RFP-KDEL fluorescence intensity in the cytoplasm near the nucleus. Image stacks with paired RFP-KDEL and GFP-tubulin fluorescence images were used for the analyses. To determine the centrosome-adjacent fluorescence measure (ratio numerator), an image within a Z-stack that bisected a single centrosome was identified based on GFP-tubulin fluorescence. The average RFP-KDEL fluorescence intensity within a 2.5 μm diameter circle immediately adjacent to the centrosome was then obtained using ImageJ. The same circle was then used to determine the average RFP-KDEL fluorescence intensity in the cytoplasm about 2 μm from the nuclear boundary in the same image (ratio denominator). Ratios were calculated at 2 min intervals, and the Z-plane of the images used was adjusted over time as needed to follow the same centrosome. For calculation of mean ER_centrosome_ / ER_cytoplasm_ during interphase and metaphase, the ratio at 40 min prior to NEB was used for interphase and the ratio at 20 min following NEB was used for metaphase; these times were determined based on staging of control cells.

### RT-qPCR

RNA was extracted from 2^nd^ instar larvae using TRIzol reagent (Invitrogen) and the Direct-zol RNA isolation kit (Zymo Research). The high capacity cDNA Reserve Transcription kit (ThermoFisher) was then used to convert RNA to cDNA in a S1000 Thermo Cycler (BioRad). RT-qPCR was carried out in a StepOnePlus cycler and detection system (Applied Biosystems). Each reaction consisted of triplicate samples containing iTaq Universal Probes Supermix (BioRad), TaqMan primers 5′ end labeled with 6-carboxyfluorescein (FAM; ThermoFisher), and cDNA diluted per manufacturer’s instructions. Pre-validated Taqman primers targeted to *Drosophila Dhc64C*, *Klp61F*, *ncd*, and *Rpl140* (housekeeping gene) were used. For quantification, triplicate cycle threshold (Ct) measures were averaged and normalized to an average Ct value for RPL140 to calculate the ΔCt. The Δ(ΔCt was then determined by subtracting the ΔCt value obtained from *GAL4* RNAi animals (non-targeting control) from the ΔCt value obtained from the experimental RNAi animals. Fold changes were expressed as 2^−Δ(ΔCt)^.

## Acknowledgements

We thank Dr. Gregory P. Mueller for critical reading of the manuscript. This work was supported by Department of Defense start-up funds to J.T.S.

## Author Contributions

D.K. designed and carried out experiments, and J.T.S. designed experiments and wrote the manuscript.

## Additional Information

The authors declare no competing interests.

