## Supplemental Figures S1-S4, Supplementary Video Legends for "A novel, dynein-independent mechanism focuses the endoplasmic reticulum around spindle poles in dividing *Drosophila* spermatocytes"

### Supplementary Materials

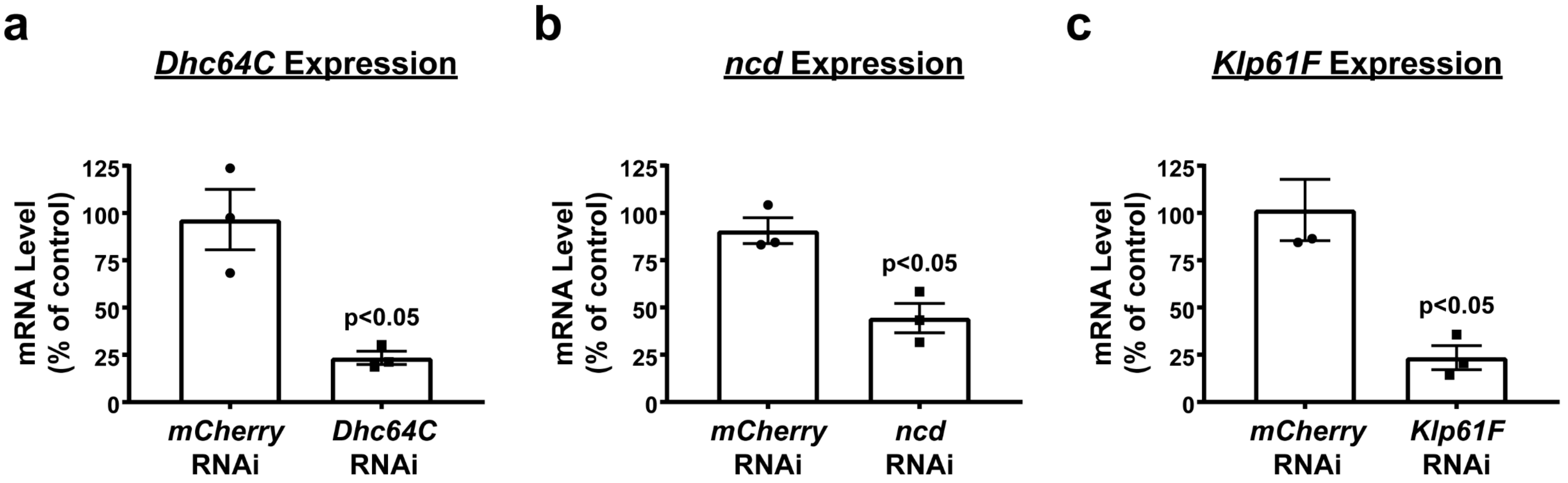


**Figure S1** Karabasheva et al

**<u>Supplementary Figure S1.</u> Suppression of D*hc64C*, K*lp61F*, and *ncd* transcript levels by RNAi.** *mCherry* (non-targeting control), *Dhc64C*, *Klp61F*, and *ncd* RNAi were expressed ubiquitously in whole animals using the *da-GAL4* driver, and total RNA was isolated from second instar larvae and analyzed by qPCR. Second instar larvae were used because *Dhc64C* and *Klp61F* RNAi caused lethality at later stages. Transcript levels were quantified using the Δ(ΔCt) method (see Methods), and values were normalized against a second, independent non-targeting control RNAi. Data are expressed as percent ± SEM from three independent experiments. Indicated p-values are for each experimental RNAi compared to mCherry (t-test).

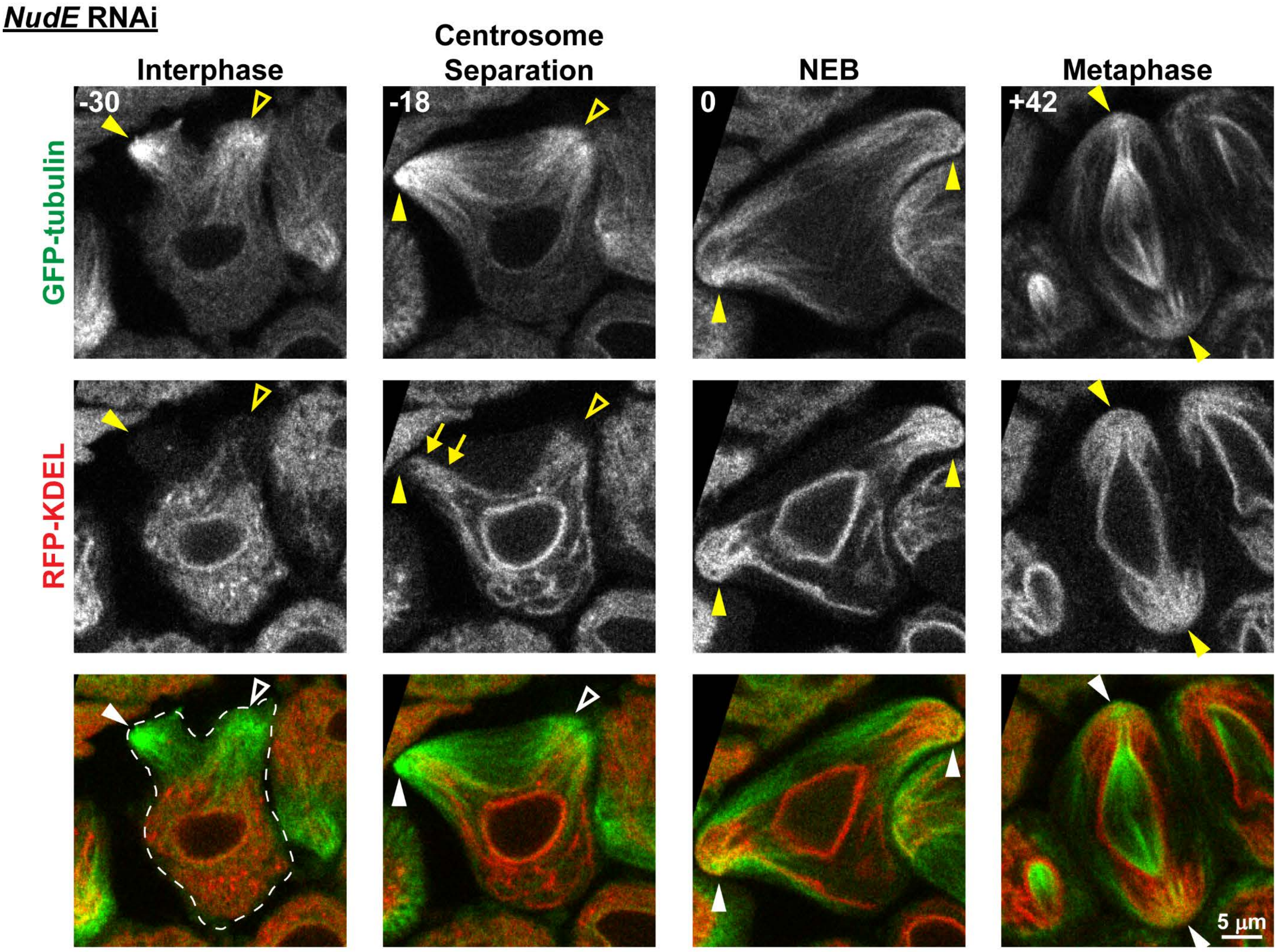


**Figure S2** Karabasheva et al

**Supplementary Figure S2. *NudE* suppression results in similar spindle and ER defects as *Dhc64c*.** Representative images of a *NudE* RNAi expressing spermatocyte with GFP-tubulin (green) and RFP-KDEL (red) from late interphase (-30 min) through metaphase (+42 min) of meiosis (times relative to NEB). Filled arrowheads indicate visible centrosomes, and open arrowheads indicate the approximate location of centrosomes that are out of the focal plane. Note the lack of RFP-KDEL fluorescence around the cortical centrosomes in interphase. Arrows in the "Centrosome Separation" image (-18 min) point to RFP-KDEL – labeled ER that is being drawn to the cortical centrosomes along the astral MTs.

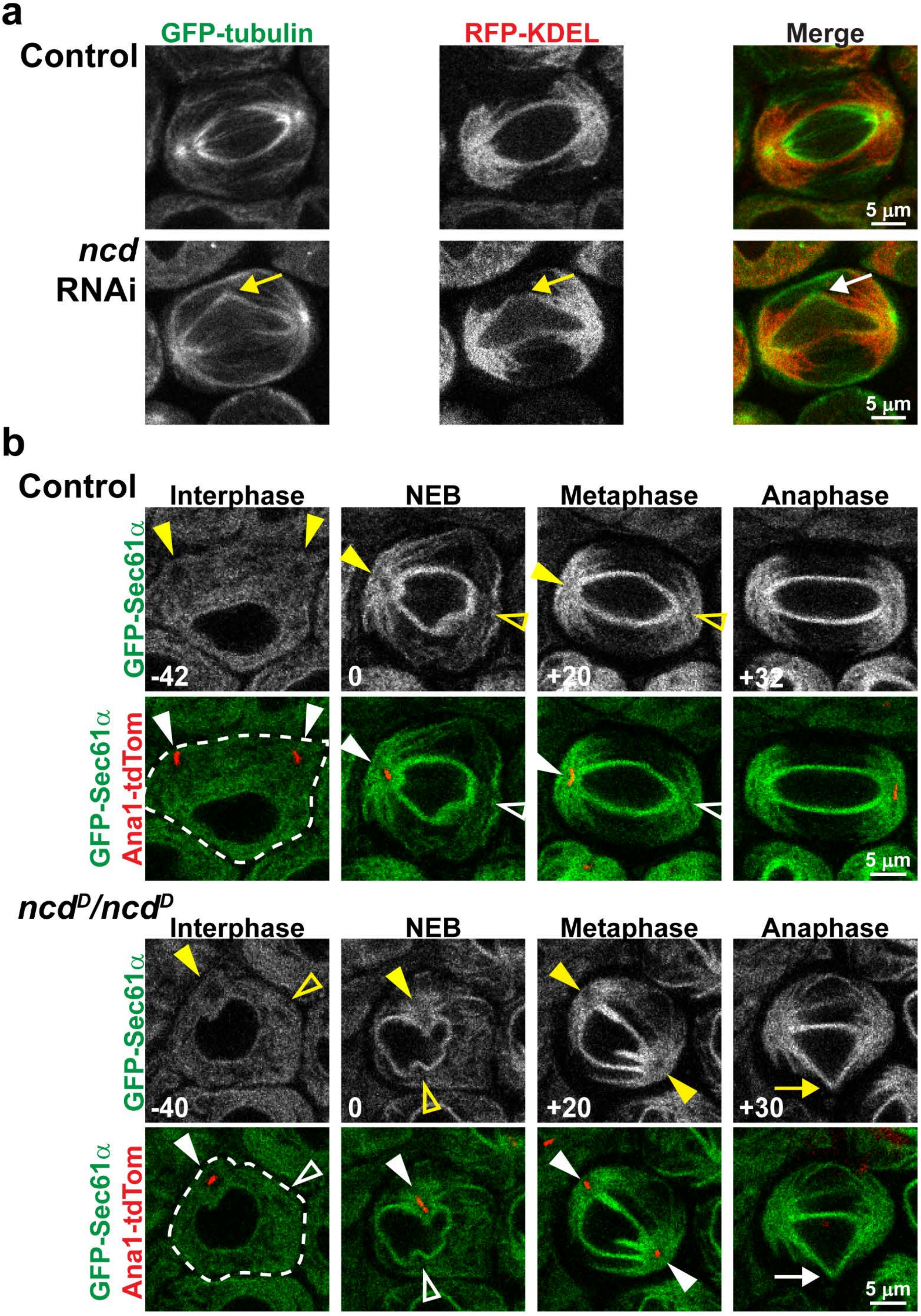


**Figure S3** Karabasheva et al

**<u>Supplementary Figure S3.</u> The *ncd*$^{D}$ mutant allele phenocopies *ncd* RNAi.** a) Representative images of a control spermatocyte (upper panel) and the *ncd* RNAi spermatocyte shown in Figure 5a (lower panel), just prior to anaphase onset. The arrow points to the region of the spindle in the *ncd* RNAi spermatocyte that is deflected to the cell cortex. b) Representative images of a control (upper panel) and *ncd*$^{D}$ / *ncd*$^{D}$ (lower panel) spermatocyte expressing GFP-Sec61α to label the ER (green) and Ana1-tdTomato (red) from late interphase through anaphase. Filled arrowheads indicate visible centrosomes, and open arrowheads indicate the approximate location of centrosomes that are out of the focal plane. The arrow in the *ncd*$^{D}$ / *ncd*$^{D}$ "Anaphase" image points to the deflected parafusorial membranes that surround the spindle, indicative of deflection of the spindle itself.

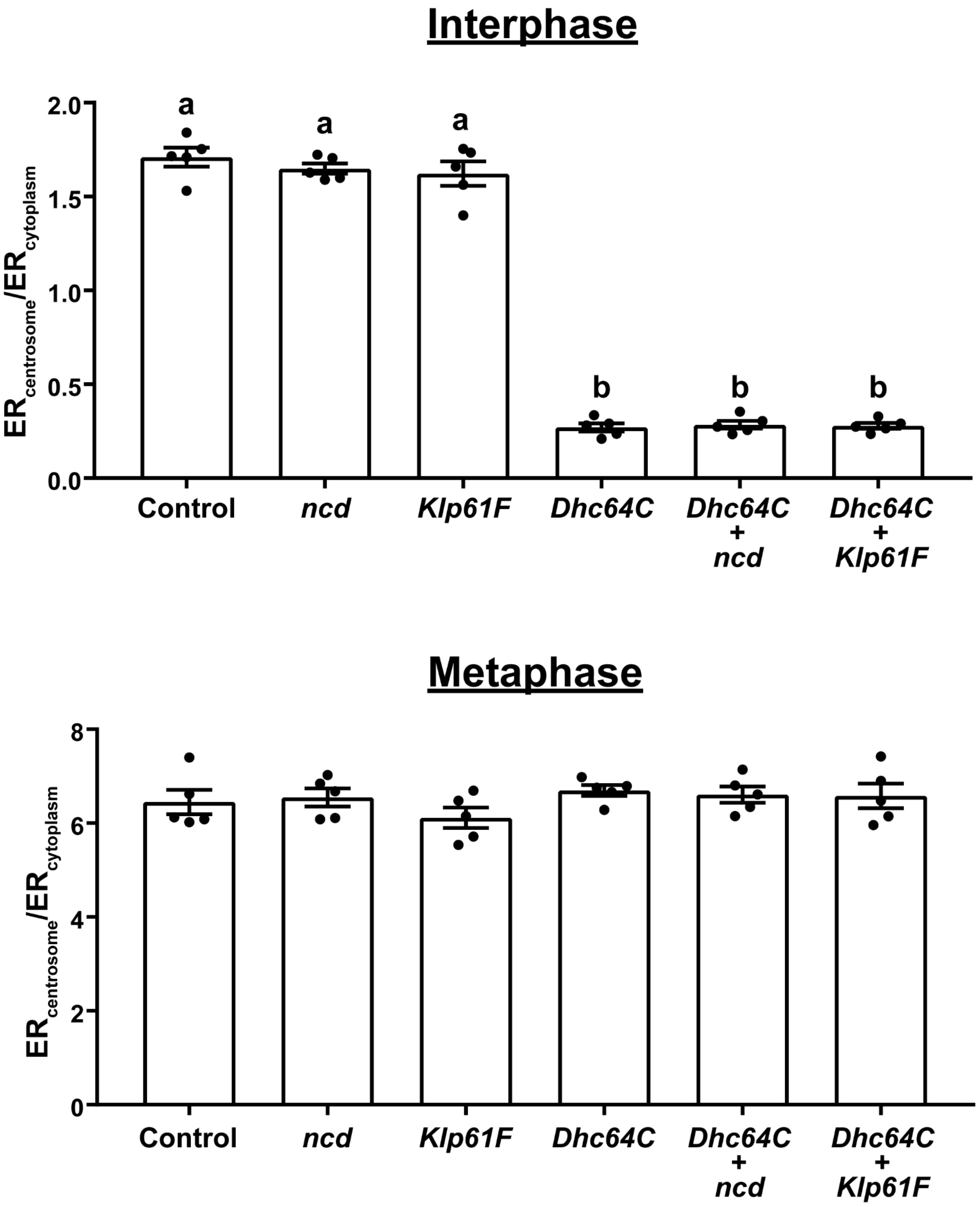


**Figure S4** Karabasheva et al

**<u>Supplementary Figure S4.</u> $ER_{centrosome}$ / $ER_{cytoplasm}$ measurements from *ncd* and *Klp61F* RNAi spermatocytes.** $ER_{centrosome}$ / $ER_{cytoplasm}$ was measured from five cells expressed RFP-KDEL and GFP-tubulin for each of the RNAi conditions indicated. Mean ± SEM is shown for interphase (measured at NEB -40 min; upper panel) and metaphase (NEB +20 min; lower panel). Different letters above the bars indicate statistical significance (one-way ANOVA with Tukey's multiple comparisons test; $p < 0.0001$).

**Supplementary Videos**

**Supplementary Video 1.** Full timelapse of the control spermatocyte shown in Figure 2a. GFP-tubulin is green and RFP-KDEL is red in the merged panel on the right. Note the centrosomes at the upper cortex of the cell at the start of the timelapse, and the concentration of RFP-KDEL fluorescence around the centrosomes as they separate and engage with the nuclear envelope. Images were acquired every 2 minutes, and the video playback rate is 2 frames/second. The scalebar represents 5 μm.

**Supplementary Video 2.** Full timelapse of the control spermatocyte shown in Figure 3a. Ana1-tdTomato and H2A-RFP are red and GFP-PDI is green in the merged panel on the right. Note that only a single centrosome, indicated by ana1-tdTomato fluorescence, is visible in the focal plane shown. Images were acquired every 60 seconds, and the video playback rate is 4 frames/second. The scalebar represents 5 μm.

**Supplementary Video 3.** Full timelapse of the *Dhc64C* RNAi spermatocyte shown in Figure 2b. GFP-tubulin is green and RFP-KDEL is red in the merged panel on the right. Note the centrosomes at the upper cortex of the cell at the start of the timelapse, and the movement of ER, indicated by of RFP-KDEL fluorescence, toward the cortical centrosomes as the centrosomes begin to separate. Images were acquired every 2 minutes, and the video playback rate is 2 frames/second. The scalebar represents 5 μm.

**Supplementary Video 4.** Full timelapse of the *Dhc64C* RNAi spermatocyte shown in Figure 3b. Ana1-tdTomato and H2A-RFP are red and GFP-PDI is green in the merged panel on the right. Only a single centrosome, indicated by ana1-tdTomato fluorescence, is visible in the focal plane shown. Note the sudden movement of the ER (GFP-PDI) toward the cortical centrosome. Images were acquired every 60 seconds, and the video playback rate is 4 frames/second. The scalebar represents 5 μm.
